# Molecular epidemiology of *Theileria parva* in Eastern Democratic Republic of Congo: Implications to the introduction of the Muguga Cocktail vaccine for East Coast Fever

**DOI:** 10.1101/2025.05.07.652749

**Authors:** Walter Muleya, Simbuwa Mulonga, Victor Mbao, Kalume Moise Kasereka, Kambale Mbusa Héritier, Boniface Namangala, Jeremy Salt, Anthony Jim Musoke, David Kalenzi Atuhaire

**Affiliations:** Department of Biomedical Sciences, School of Veterinary Medicine, University of Zambia, Lusaka, Zambia; International Development Research Centre, Eastern and Southern Africa Regional Office, Nairobi, Kenya; Faculty of Veterinary Medicine, The Catholic University of Graben, Butembo, DR Congo; Department of Paraclinical Studies, School of Veterinary Medicine, University of Zambia, Lusaka, Zambia; Global Alliance for Livestock Veterinary Medicines, Pentlands Science Park, Bush Loan, Penicuik Edinburgh, Scotland. (Previous address); The Vaccine Group, Ltd, Sandwich, England, United Kingdom. (Current address); LMK Medical laboratories and Consultancies, Kampala, Uganda; Centre for Ticks and Tick-Borne Diseases, Lilongwe, Malawi. (Previous address); College of Veterinary Medicine Animal Resources and Biosecurity, Makerere University, Kampala, Uganda. (Current address)

**Keywords:** *Theileria parva*, DRC, Immunization, Genetic diversity, Field challenge trial

## Abstract

East Coast Fever (ECF) is one of the most economically important tick-borne diseases of cattle in Eastern, Central, and Southern Africa, caused by the intracellular protozoan parasite *Theileria parva* (*T. parva*). This study investigated the prevalence and genetic diversity of *T. parva* populations in the eastern Democratic Republic of Congo (DRC) to inform immunization strategies against ECF. By employing PCR and DNA sequencing techniques, in conjunction with surveys assessing Knowledge and Practices, as well as conducting an immunization and field challenge trial, the findings suggest the existence of *Rhipicephalus appendiculatus* ticks in the provinces of South-Kivu, North Kivu, and Ituri. This underscores a possible threat of disease transmission in these areas. Molecular analyses uncovered diverse *T. parva* populations with varying antigenic profiles, challenging the assumption of uniformity despite Muguga cocktail-like appearances. Moreover, phylogenetic analyses suggested limited similarity between *T. parva* populations and the Muguga cocktail vaccine. Microsatellite analysis and field challenge trials supported the notion of multiple populations, highlighting the current vaccine’s limitations against field strains. This study has the potential to significantly contribute to understanding *T. parva* dynamics in the region, emphasizing the complexities of vaccine strain selection and stressing the importance of continuous monitoring and adaptive control strategies in the face of evolving parasite populations.

## Introduction

East Coast Fever (ECF) is one of the most economically significant tick-borne diseases affecting cattle in Eastern, Central, and Southern Africa, caused by the intracellular protozoan parasite *Theileria parva*; (1,2). The transmission of this disease occurs through the three-host tick, *Rhipicephalus appendiculatus* (3), predominantly parasitizing cattle (4). In the Democratic Republic of the Congo (DRC), ticks belonging to the Rhipicephalus genus are notably found to be most abundant in cattle (5). Research conducted in North Kivu, a province in the eastern part of DRC, focused on identifying tick species and their correlation with the seroprevalence of *T. parva* in cattle raised under an extensive farming system, revealed dominance of *R. appendiculatus* in the area, suggestive of a potential state of endemicity to *T. parva* infection in the area (6).

The life cycle of *T. parva* is intricate, involving obligatory developmental stages in both mammalian and vector hosts (7). Cattle become infected through the inoculation of sporozoite forms present in tick saliva. The disease significantly affects exotic and crossbred cattle, along with indigenous calves under six months old, resulting in elevated mortality rates. In endemic areas, chronic ECF manifests with less quantifiable effects such as reduced growth, diminished milk production, poor weight gain, low fertility rates, paralysis, and vulnerability to secondary attacks from other parasites (8). Consequently, ECF stands as a primary motivator for tick control measures in numerous African countries.

Control of ECF involves the use of a combined system of acaricides, chemotherapy, livestock movement and Infection and Treatment Method (ITM). The utilization of acaricides remains a widespread method of tick control, albeit obstacles such as elevated expenses, emergence of tick resistance and insufficient dipping facilities. Despite these challenges, a considerable number of farmers resort to acaricides for ECF control even though the most effective preventive approach is the infection and treatment method (ITM). ITM involves the simultaneous inoculation of a live, potentially lethal dose of *T. parva* sporozoites and a long-acting oxytetracycline (9). While ITM is stock-specific, combinations of stocks, such as the widely used ‘Muguga cocktail’ (10) comprising *T. parva* Muguga, Kiambu 5, and Serengeti-transformed stocks, provide broad protection. Despite its efficacy, ITM adoption faces challenges due to cold chain difficulties and concerns about vaccine strains establishing in resident tick populations and intermingling with local parasite genotypes (11).

The effectiveness of the ITM in providing protection is contingent on the selection of an immunizing stock, emphasizing the need for characterizing local strains in the Eastern Democratic Republic of Congo. Characterization is essential to maximize cross-immunization possibilities. While there is no direct correlation between molecular profiling and immunogenicity, characterizing local strains helps narrow down the parasite stocks or isolates for cross-immunization trials, thereby reducing the cost of introducing immunization. For comparative purposes, different and well-characterized strains are often included in a single vaccine, exemplified by the success of the Muguga Cocktail in immunization trials.

The sequencing of *T. parva* CTL antigens has emerged as a valuable tool for assessing the diversity of *T. parva* in endemic areas (12–15). In addition, the identification of polymorphic minisatellites and microsatellite markers in the *T. parva* genome allows direct genotyping of isolates from blood samples using specific primers (16–18). These locus-specific markers, based on variable number tandem repeats (VNTRs) in non-coding regions, serve as valuable tools for studying population structure due to their technical simplicity in analysing repeat motif copy number variation. The utilization of minisatellites and microsatellites in *T. parva* genotyping has unveiled diverse population structures (17–20).

Numerous studies highlight substantial diversity among *T. parva* isolates, with observed genetic exchange when multiple stocks infect the tick vector (13,16–23).

In the DRC, communities in the eastern regions heavily depend on livestock for their livelihoods and the presence of theileriosis poses a significant impediment to the growth and enhancement of the livestock sector. Consequently, implementing effective control measures, particularly through immunization, is crucial, given its cost-effectiveness compared to chemotherapy. Thus, with the recognition of the importance of genetic exchange in *T. parva* populations, this study aimed to assess the seroprevalence and conduct population genetic analysis for potential vaccine candidates against ECF in Eastern DRC. Utilizing sequence and microsatellite analysis, coupled with an immunisation and field challenge trial, this approach sought to identify effective immunization strategies.

## Materials and Methods

### Study sites

Initially, to determine the prevalence and presence of *T. parva*, a cross-sectional study was conducted in September 2012, covering three provinces in the DRC: North Kivu, South Kivu, and Ituri, each comprising three agro-ecological zones—low altitude (< 1200m), medium altitude (1200 to 1800m), and high altitude (> 1800m) (Fig.1). The study included a total of 31 villages (North Kivu = 12, South Kivu = 8, and Ituri = 11) distributed across the provinces (Fig.1). Territories within each province were carefully selected based on their concentration of the most significant cattle population and their suitability as an agro-ecological zone for the *R. appendiculatus* tick. Specifically, two territories were chosen in North Kivu (Beni and Lubero), three in South Kivu (Kabare, Walungu, and Uvira), and two in Ituri (Irumu and Mambasa). Due to wartime occupation during the sampling period, some villages in Ituri experienced limited dispersion in terms of latitude and longitude. Consequently, certain territories such as Walikale and Rutshuru in North Kivu (with indigenous cattle populations) and Mwenga in South Kivu (housing the main wild animals) were excluded.

**Fig. 1:**
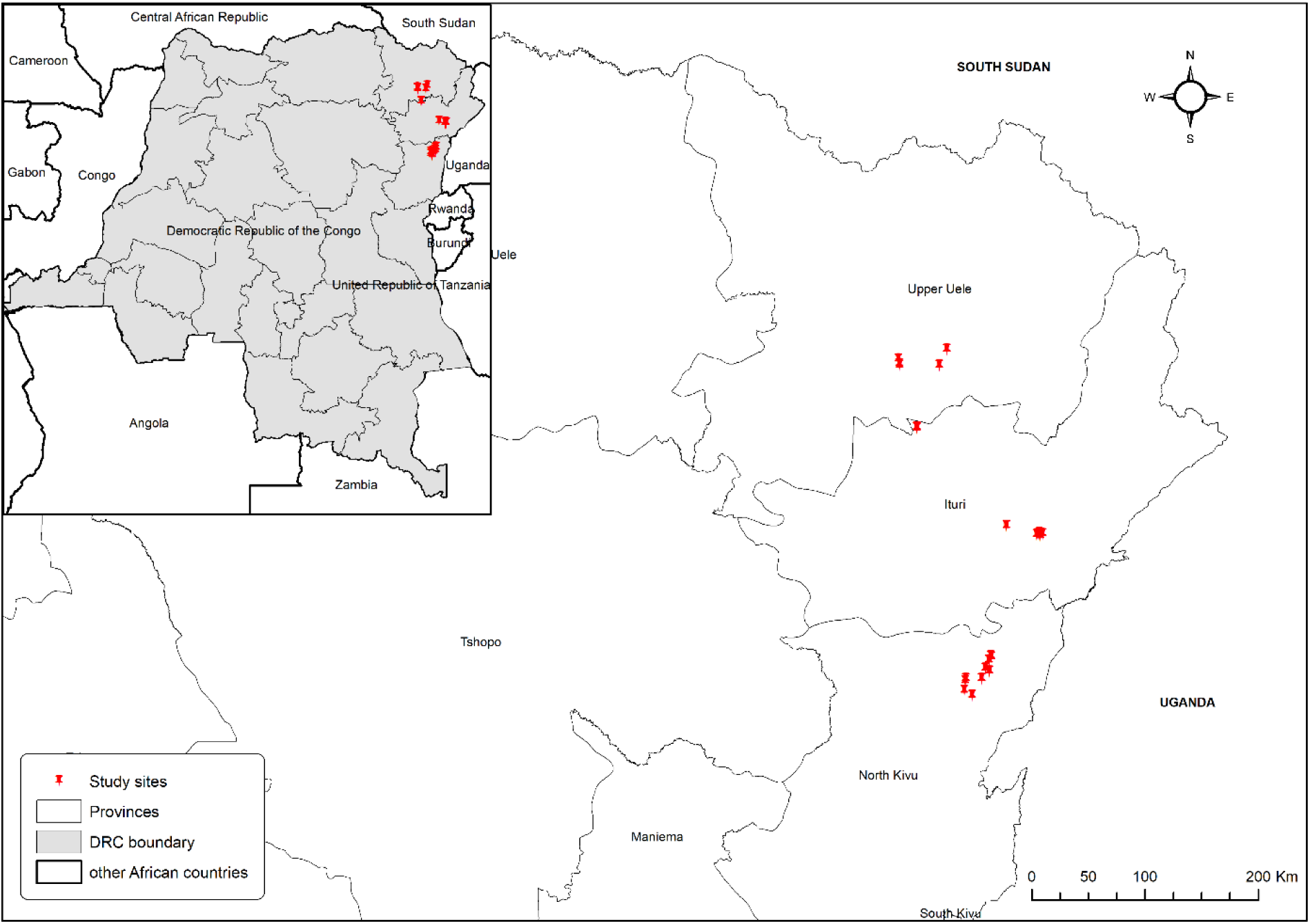
Map of DRC showing study sites

### Sample collection

Ticks were collected randomly from cattle using small forceps starting from the head towards the tail and placed in a petri dish. Care was taken to avoid decapitulation and shedding of the legs. The ticks were then transferred to plastic bottles with holes made in the cap for aeration. Blood samples were collected from cattle through jugular venipuncture using vacutainers with EDTA and plain tubes for molecular analysis and ELISA, respectively. The collected whole blood and serum were stored at -20°C and the collected ticks were identified at the Department of Parasitology and Parasitic Diseases, Faculty of Veterinary Medicine, Catholic University of Graben, Butembo, DRC. Subsequently, they were transported to the Center for Ticks and Tick-borne Diseases (CTTBD) in Lilongwe, Malawi, for *T. parva* ELISA and PCR screening.

### ELISA and p104 *T. parva* diagnostic PCR

The PIM ELISA was used for *T. parva* screening of serum samples collected according to the protocol described by (24). With regards to PCR screening, DNA was purified from cattle blood samples and spotted FTA filter papers (Whatman Bio-Science) as described previously (16). Field samples were screened for *T. parva* DNA by using *T. parva* specific p104 gene primers as described previously (25).

### Questionnaire survey

Fifteen farmers were administered a pre-formed questionnaire to gather information on farm management and challenges encountered. The survey staff assisted in completing the questionnaire.

### Sentinel study sites and sampling

To determine which farm/area could be used for field immunization challenge, sentinel cattle (n=5) were placed at Kabasha village, Beni territory, North Kivu Province in November 2016. Blood on Whatman No. 4 filter paper was collected and shipped to the CTTBD in Lilongwe for DNA extraction and PCR analysis. For further characterization, DNA was sent to the School of Veterinary Medicine, University of Zambia (UNZA), Lusaka, Zambia.

### Field exposure: Immunization and Challenge trial

To select suitable animals for a *T. parva* immunization/challenge trial, sampling involved 150 cattle from 15 farms in the highland area of Lubero and Beni territories, North Kivu province, DRC, (longitude 29° 12′ to 29° 17 E and latitude 0° 8 N to 0°40 S) with year-round rainfall. Blood on filter paper was sent to CTTBD, Lilongwe, for PCR analysis. Subsequently, 100 negative cattle were purchased, confined at UCG/Butembo farm, and underwent strict tick control using SUPONA EXTRA (1/2000) to prevent premature parasite challenge. After acclimatization, the animals were divided into two groups, A (n=50) and B (n=50). Group A received 1ml of the Muguga cocktail vaccine and long-acting Oxytetracycline simultaneously, while Group B served as non-immunized controls. All animals were closely monitored for 35 days under strict tick control. Immunized animals were bled on Days 0, 21, 28, and 35 post-immunization, while the non-immunized control group was bled on Days 28 and 35. The sera were stored at -20°C at the Faculty of Veterinary Medicine at UCG/Butembo, DRC, before shipping to ILRI/Kenya for PIM ELISA analysis (24).

Before exposing both groups to tick/parasite challenge, animals underwent thorough daily washing for three consecutive days (Days 33, 34, and 35 post-immunization) to eliminate residual acaricide, facilitating easy and high tick attachment during exposure. On Day 35 post-immunization, all animals were field-exposed in Kabasha village, grazing freely. Animals with severe reactions to immunization were excluded, receiving treatment or euthanized if needed.

Throughout the exposure period, animals received no tick control or ECF treatment but were treated for other diseases. Daily temperature monitoring occurred, and if an animal exhibited a high temperature (> 39.5°C) for two consecutive days, a blood smear was checked for hemoparasites. Animals with enlarged lymph nodes underwent biopsy smears stained with Giemsa. Blood samples were further collected from animals that succumbed to ECF during the exposure period and sent to the University of Zambia, School of Veterinary Medicine for molecular characterization.

### p67, Tp1 and Tp2 PCR and cycle sequencing

The *T. parva* p67, Tp1, and Tp2 genes were amplified using the Amplitaq Gold PCR kit (Invitrogen, CA) with primers designed by (26). Briefly, in a 20 μL reaction, consisting of 10 ng template DNA, 10 μL 2x Amplitaq Gold master mix, 0.5 μM of each primer, and distilled water, the amplification cycles were as follows: denaturation at 95°C for 10 minutes, followed by 40 cycles at 96°C for 30 seconds, 55°C (p67) 50°C (Tp1) and 53°C (Tp2) for 45 seconds, and 68°C for 60 seconds, with a final extension step at 72°C for 5 minutes. PCR products were visualized on 1.5% ethidium bromide-coated agarose gels, with expected product sizes of 906 bp, 432 bp, and 525 bp for p67, Tp1, and Tp2, respectively.

The obtained PCR products underwent purification using the Monofas purification kit (GL Sciences, Japan), following the manufacturer’s instructions. Subsequently, cycle sequencing was carried out using the Big Dye Terminator v3.1 cycle sequencing kit (Life Technologies, Applied Biosystems), following the manufacturer’s instructions. The resulting cycle sequencing products were purified to remove excess labeled dNTPs, buffers, and enzymes through the ethanol precipitation method followed by denaturation. Finally, capillary electrophoresis was performed on the ABI 3500 genetic analyzer (Life Technologies).

### Microsatellite and minisatellite genotyping

The genotyping markers employed for field samples can be found in **Additional File 1**. Fluorescent dye-labeled forward primers and previously specified annealing temperatures (13) were used. PCR mix and reaction conditions followed established protocols (13). After amplification, PCR products were observed on a 1.5% agarose gel coated with ethidium bromide. Successful PCR products were denatured and electrophoresed using the ABI Seqstudio genetic analyzer (Life Technologies). DNA fragment sizes from control, immunized, sentinel and vaccine strains (Muguga, Kiambu, Serengeti, and Chitongo vaccine isolates) were determined using GeneMapper software ver. 5 (Applied Biosystem, Waltham, Massachusetts, USA). The software scored peaks with the highest area as the most dominant allele, and based on this, a Multi-locus genotype (MLG) was constructed to represent the most dominant genotype within each sample.

### Data analysis

#### ELISA and Sequence analysis

The OD values were expressed as percent positivity (PP) relative to a reference strong-positive control serum. Any test serum with a PP value of 20 or above was considered positive. For nucleotide sequence verification, Blast analysis on the NCBI website was employed. The sequences obtained were assembled and edited using Genetyx ver. 12. Alignment with Muguga, Kiambu 5, Serengeti transformed, and Chitongo vaccine strain reference sequences, along with other East African references, was performed using Clustal W1.6. A fasta file of this alignment was generated and converted to a MEGA file format for constructing phylogenetic trees with MEGA ver. 6 (27). Phylogenetic trees were constructed using 1000 bootstrap replicates. Multiple sequence alignments of amino acid sequences, translated from nucleotide alignments, were conducted to analyze CTL epitopes on both Tp1 and Tp2. DNA polymorphisms for each gene were calculated using DnaSP ver. 5 (28). The single likelihood ancestor counting method with the F81 model and a confidence level of 0.05 on the Data Monkey website (http://www.datamonkey.org) was used to calculate the mean ratio of non-synonymous substitutions vs. synonymous substitutions (dN/dS) per site (29,30). The distribution of genetic variation and population differentiation among sequences were investigated through AMOVA performed using GenAIEx6 (31).

Network ver. 10 was utilized to assess haplotype similarities between Tp1 and Tp2 nucleotide sequences of the vaccine stocks and field samples (http://fluxus-engineering.com/). All nucleotide sequences obtained in this study have been deposited in the DNA Data Bank of Japan (DDBJ) with designated accession numbers LC863893-LC863940.

#### Microsatellite analysis

Initially, microsatellite analysis was employed using the microsatellite toolkit (http://animalgenomics.ucd.ie/sdepark/ms-toolkit/) to assess the level of similarity within the MLG. The results were visualized through allele frequency distribution and Principal Component Analysis (PCA) created using GenAIEx6 (31). Population sub-structuring was quantified using the FSTAT computer package version 2.9.3.2 (https://www2.unil.ch/popgen/softwares/fstat.htm).

To test the null hypothesis of panmixia and linkage equilibrium, LIAN (Haubold and Hudson, 2000) was employed. This involved calculating the standardized index of association, the variance of pairwise differences (VD), the variance of differences required for panmixia (VE), and L, representing the 95% confidence interval for VD. Negative or close-to-zero values of the index of association suggest panmixia (random mating), while positive values significantly greater than zero indicate non-panmixia (nonrandom mating). The rejection of the null hypothesis of panmixia is indicated when VD is greater than the L value, signifying linkage disequilibrium (LD). Conversely, acceptance of the null hypothesis of panmixia occurs when the calculated VD is less than the L value, indicating linkage equilibrium (LE).

## Results

### Characteristics of cattle in the study area

Three variables were taken into account for sampling: (i) age group, (ii) sex, and (iii) agro-ecological zone. A minimum of 250 animals were sampled from each province, resulting in a total of 756 animals. The majority of the sampled animals were adult females (> 24 months) (Table 1), reflecting the common practice of slaughtering males before they reach 12 months of age. The distribution of animals was consistent across all agro-ecological zones (AEZ).

**Table 1:** Distribution of the characteristics of cattle in the study area.

| Variables | Levels | Frequency | Percentage |
| --- | --- | --- | --- |
| Age group (Month) | < 6 | 3 | 0.4 |
|  | 6 to 12 | 106 | 14.02 |
|  | 13 to 24 | 115 | 15.21 |
|  | > 24 | 532 | 70.37 |
| Sex | Males | 194 | 25.66 |
|  | Females | 562 | 74.34 |
| Agro-ecological zone | Low altitude (<1200m) | 355 | 46.96 |
|  | Medium altitude (1200 to 1800m) | 270 | 37.71 |
|  | High altitude (> 1800m) | 131 | 17.33 |

### Distribution of ticks

A total of 12,672 ticks were gathered from 756 animals, and five tick species were identified, namely: *R. appendiculatus* (68.04%), *R. decoloratus* (18.16%), *Amblyomma variegatum* (7.87%), *R. evertsi evertsi* (5.7%), and *Hyalomma m. rufipes* (0.24%). Notably, *R. appendiculatus* emerged as the most prevalent species, constituting the majority at 68.04% (Table 2).

**Table 2:**
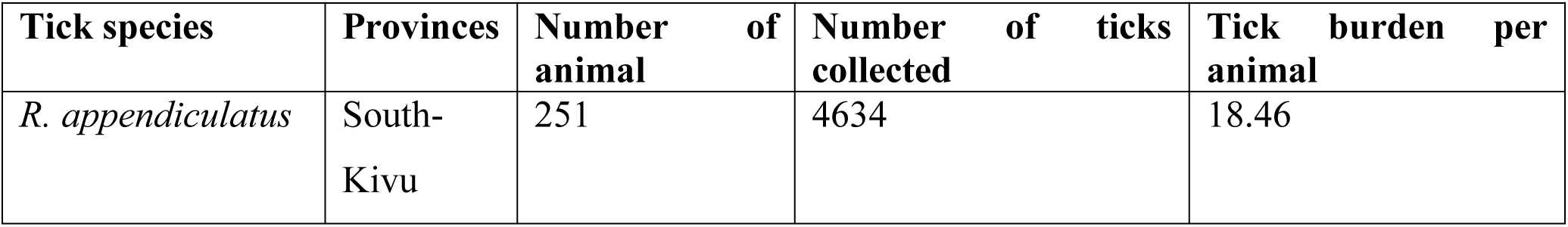
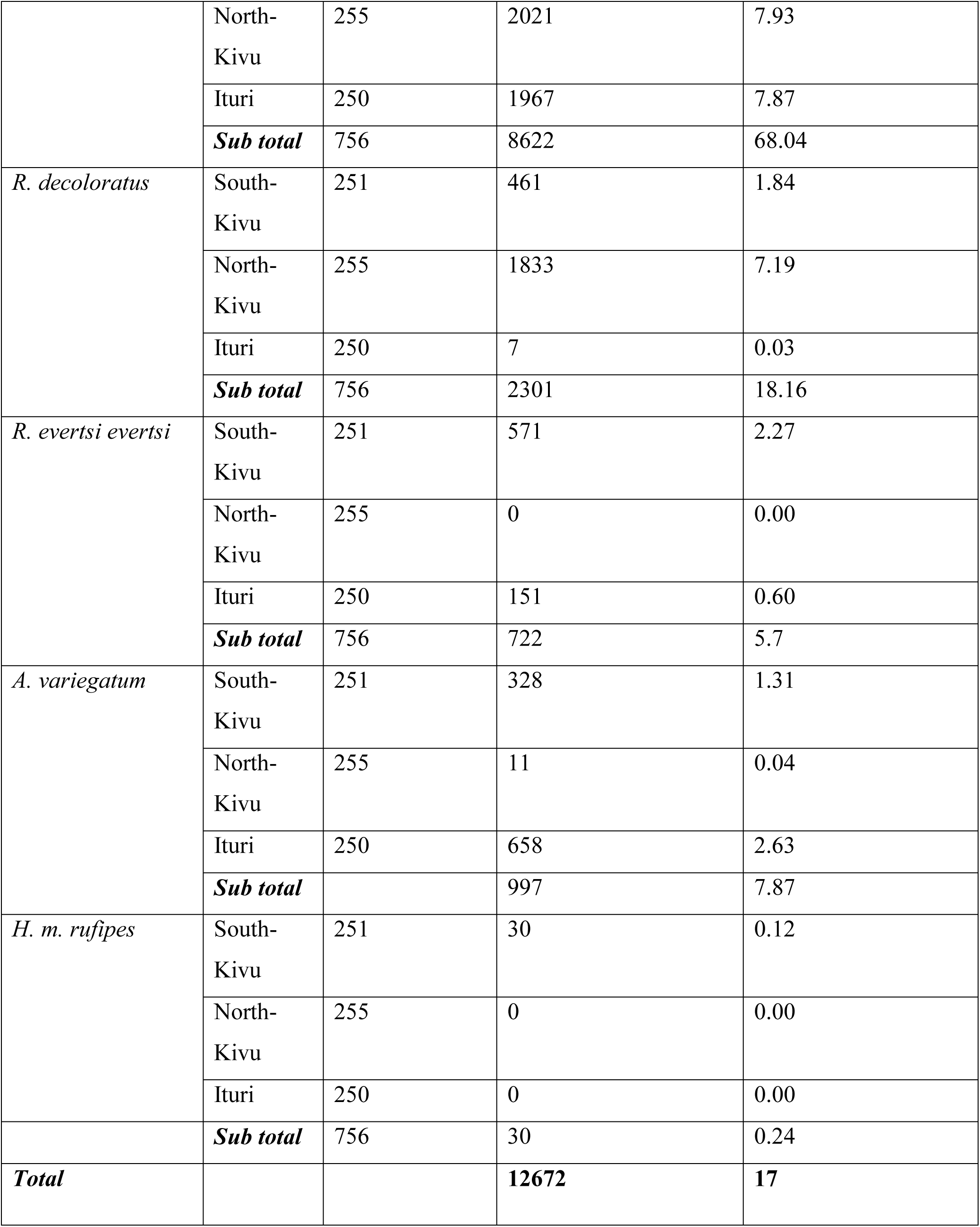
Tick populations collected from cattle in DRC’s three provinces: Ituri, North Kivu and South Kivu.

### ELISA and PCR screening for *T. parva*

Out of the 756 serum samples analyzed, 208 (27.5%) tested positive for *T. parva*, and 24 yielded inconclusive results on ELISA (Table 3). Ituri province exhibited the highest prevalence, closely followed by South Kivu. Regarding the p104 gene-specific PCR detection of *T. parva*, out of the total 208 ELISA positive samples, 182 (87.5%) tested positive (Table 4). The highest prevalence was observed in North Kivu, followed by South Kivu and Ituri provinces.

**Table 3:**
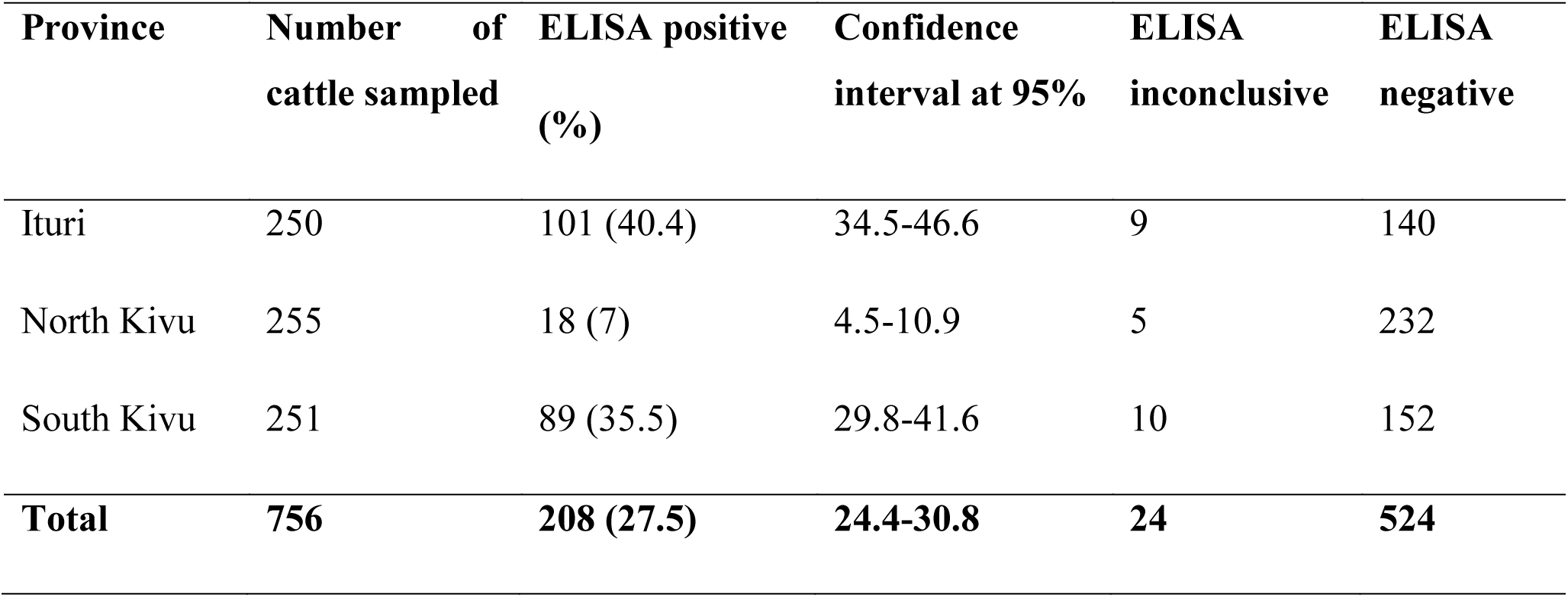
Summary of cattle *T.parva* antibodies detected using PIM ELISA.

| Province | Number of cattle sampled | ELISA positive (%) | Confidence interval at 95% | ELISA inconclusive | ELISA negative |
| --- | --- | --- | --- | --- | --- |
| Ituri | 250 | 101 (40.4) | 34.5-46.6 | 9 | 140 |
| North Kivu | 255 | 18 (7) | 4.5-10.9 | 5 | 232 |
| South Kivu | 251 | 89 (35.5) | 29.8-41.6 | 10 | 152 |
| **Total** | **756** | **208 (27.5)** | **24.4-30.8** | **24** | **524** |

**Table 4:** PCR detection of *T. parva* using p104 gene.

| Province | Number of samples | PCR positive (%) | Confidence interval at 95% | PCR negative |
| --- | --- | --- | --- | --- |
| Ituri | 101 | 86 (85.1) | 76.8-90.8 | 15 |
| North Kivu | 18 | 18 (100) | - | 0 |
| South Kivu | 89 | 78 (87.6) | 79-93 | 11 |
| **Total** | **208** | **182 (87.5)** | **82.3-91.3** | **26** |

### Responses of farmers to the questionnaire

The survey participants consisted predominantly of young farmers, aged between 20 and 36. Among them, 40% had no formal education, 13% had completed primary education, while 47% had discontinued their secondary education. The number of cattle per farm varied from 15 to 150, with an average of 48 head of cattle. All the cattle were crossbreeds, a mix of local breeds (Ankole or Ndama) and exotic breeds (Brown Swiss or Fresians).

The primary reported ailment was ECF, although some farmers expressed concerns about the thinness of their animals, potentially indicating malnutrition issues. Notably, none of the farmers had vaccinated their animals against ECF. The primary method for ECF control relied on acaricide application through hand spraying, conducted either weekly or bi-monthly. Despite these efforts, farmers observed a lack of effectiveness in acaricide application, leading to losses from ECF. This suggests that the current control method may be inadequate.

All farmers privately purchased their acaricides from Agrovet shops, as there were no government extension services available. Veterinary services were exclusively provided by private veterinarians. Most farmers had not undergone any agricultural training, highlighting a general lack of awareness regarding tick and tick-borne disease control. Only one farmer demonstrated awareness of vaccination as a potential method for controlling tick-borne diseases.

### ECF Reactions of sentinel cattle

Over the 20-day exposure period, the sentinel cattle placed at Kabasha village succumbed to a severe form of ECF, resulting in the death of four animals. Postmortem examinations revealed significant pulmonary edema and hemorrhages in the abomasum and small intestines. Schizonts were also identified in impression smears of the spleen, liver, and lymph nodes. The fifth animal experienced moderate to severe reactions but ultimately recovered (Table 5). Based on these findings, Kabasha village was selected as a suitable location for the immunization/challenge trial.

**Table 5:** ECF Reactions of the 5 Sentinel Cattle placed at Kabasha Village.

| **Animal ID Number** | **Days to Temperature** | **Duration of Temp** | **Day to Schizonts** | **Duration of Schizonts** | **Day to Piroplasms** | **Duration of Piroplasms** | **Time to euthanasia** | **Classification of Reactions\*** |
| --- | --- | --- | --- | --- | --- | --- | --- | --- |
| UC-16-8 | 23 | 17 | 23 | 11 | 20 | 14 | 34 | SR |
| UC-18-24 | 13 | 12 | 15 | 9 | 14 | 11 | 24 | SR |
| UC-20-19 | 15 | 9 | 16 | 7 | 15 | 8 | 23 | SR |
| UC-21-12 | - | - | 30 | 13 | 35 | ND | - | MOD |
| JO-9-11 | 23 | 7 | 25 | 4 | 21 | 8 | 29 | SR |

**NR:** Non-Reactor, **MR:** Mild Reactor, **MOD:** Moderate Reactor, **SR:** Severe Reactor.

### Immunization / Tick parasite challenge trial

Serological analysis of the 50 cattle immunized with the stabilate revealed a commendable success rate, with 94% (47 animals) demonstrating sero-conversion. In stark contrast, among the non-immunized controls, only 14% (7 cattle) exhibited cross-reactions with PIM, and 86% fell below the cut-off level, signifying successful immunization. Unfortunately, seven of the initially immunized cattle succumbed to intercurrent infections before detectable East Coast Fever (ECF) emerged, necessitating their exclusion from the analysis.

Among the remaining 43, 23 cattle succumbed to severe ECF, while 20 successfully withstood the field challenge (46.5%). Nine fell into the non-reactor category, five in the mild category, and six in the moderate category (Table 6). In the control group, all animals had signs of severe ECF and were euthanized. Postmortem lesions were pathognomonic for ECF.

**Table 6:** ECF reactions of both immunized and control animals.

| **Experimental Group** | **Number of Animals** | **NR** | **MR** | **MOD** | **SR** | **Number Surviving** | **Percent Protection** |
| --- | --- | --- | --- | --- | --- | --- | --- |
| Control | 49 | 0 | 0 | 0 | 49 | 0 | 0 |
| Immunized | 43 | 9 | 5 | 6 | 27 | 20 | 46.5 |

NR: Non-reactor, MR: Mild reactor, MOD: Moderate reactor, SR: Severe reactor.

### Sequence diversity and phylogenetic analysis of the p67, Tp1 and Tp2 genes p67 locus

The region spanning from 2946 to 3767 base pairs of the p67 gene, characterized by an exon-intron-exon structure, was sequenced in control, immunized and sentinel samples. Four p67 alleles were recognised as follows: allele 1 (with a 129 bp deletion), allele 2 (without deletion) (32), Allele 3 (with a 179 bp deletion), and Allele 4 (lacking the 179 bp deletion but sharing sequence similarity with Allele 3) (33). In all the study sequences, a consistent 129 bp deletion was observed, and upon phylogenetic analysis (Fig. 2), these sequences clustered with the reference sequences of allele 1. None of the sequences clustered with the remaining alleles (Fig. 2).

**Fig. 2:**
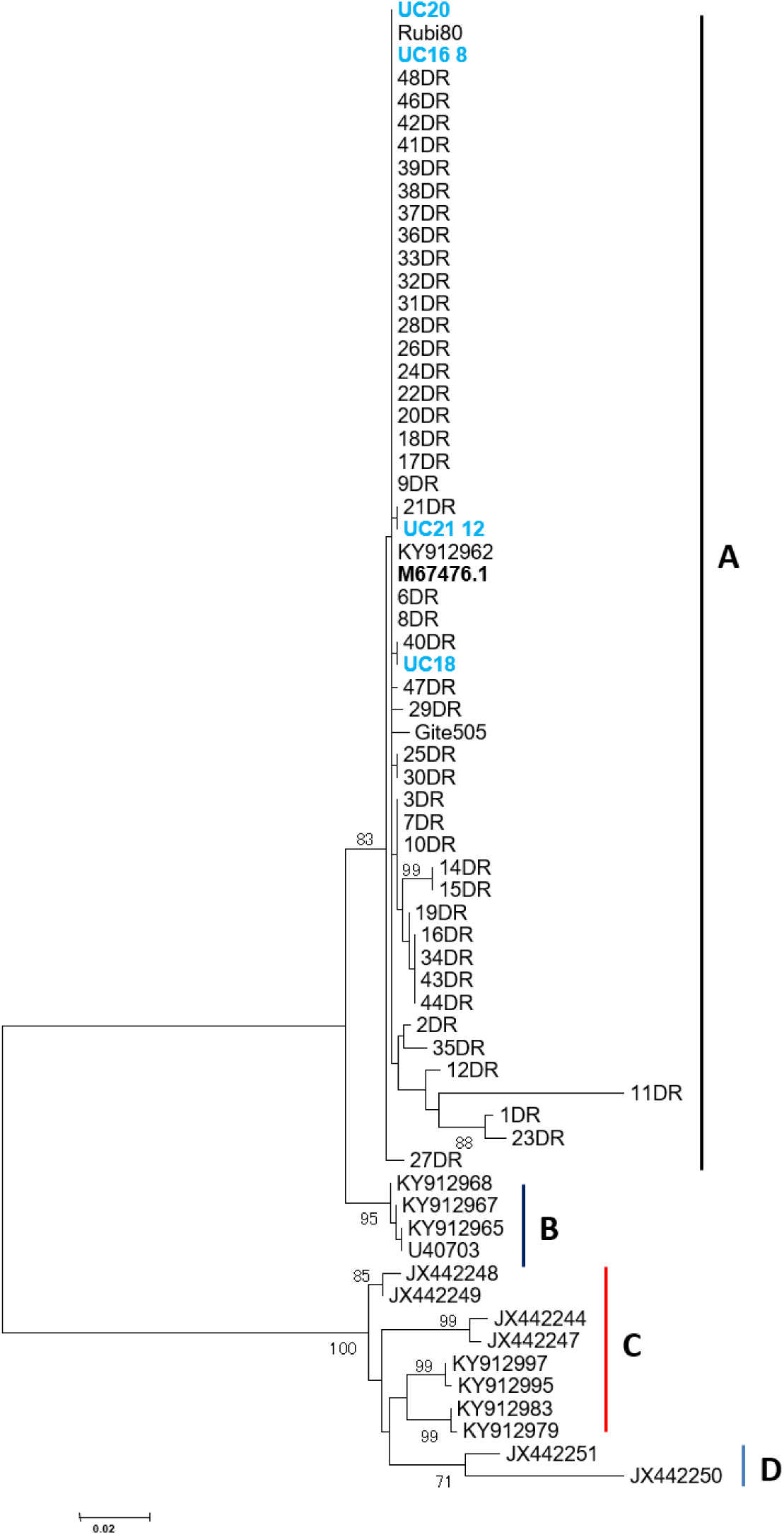
p67 phylogenetic tree based on 821 bp nucleotide sequences from sentinel and field challenge sequences from the Democratic Republic of the Congo along with reference sequences from East and Southern Africa. The tree was constructed using MEGA version 6 with 1000 bootstrap replicates as a measure of confidence.

### Tp1 locus

The Tp1 CTL gene (432 bp) of *T. parva* was sequenced from the p104 gene-positive samples. A total of 33 samples exhibited 100% amino acid sequence homology with the MC epitope. Among these, two samples (UC16, UC18) were from the sentinel group, while the remaining 31 were from the immunized (n=13) and control (n=18) groups. Three variant epitopes were observed: VGYPKVEEEML (n=1, immunized), VGYPKVKEEMI (n=4, all immunized), and VGYPKVKEEII (n=9, 5 immunized, 3 controls, and 1 sentinel). The identified epitopes in the control, immunized, and sentinel groups were largely similar to those found in the Muguga cocktail. Collectively, the samples produced a DNA polymorphism of 0.6% and a mean ratio of dN/dS of 2.15, with one positive selection site indicating positive selection.

Tp1 gene phylogenetic analysis revealed that all study sequences clustered in major cluster A. Within this cluster, samples UC16 and UC18 from sentinel cattle formed a minor cluster together with the MC in cluster AIIa, while UC20 clustered in AIa (Fig. 3). Sequences from both controls and immunized animals segregated into two minor clusters AIa and AIIb. Cluster AIIb was closely related to the Muguga cocktail strains (cluster AIIa), while cluster AIa was not (Fig. 3).

**Fig. 3:**
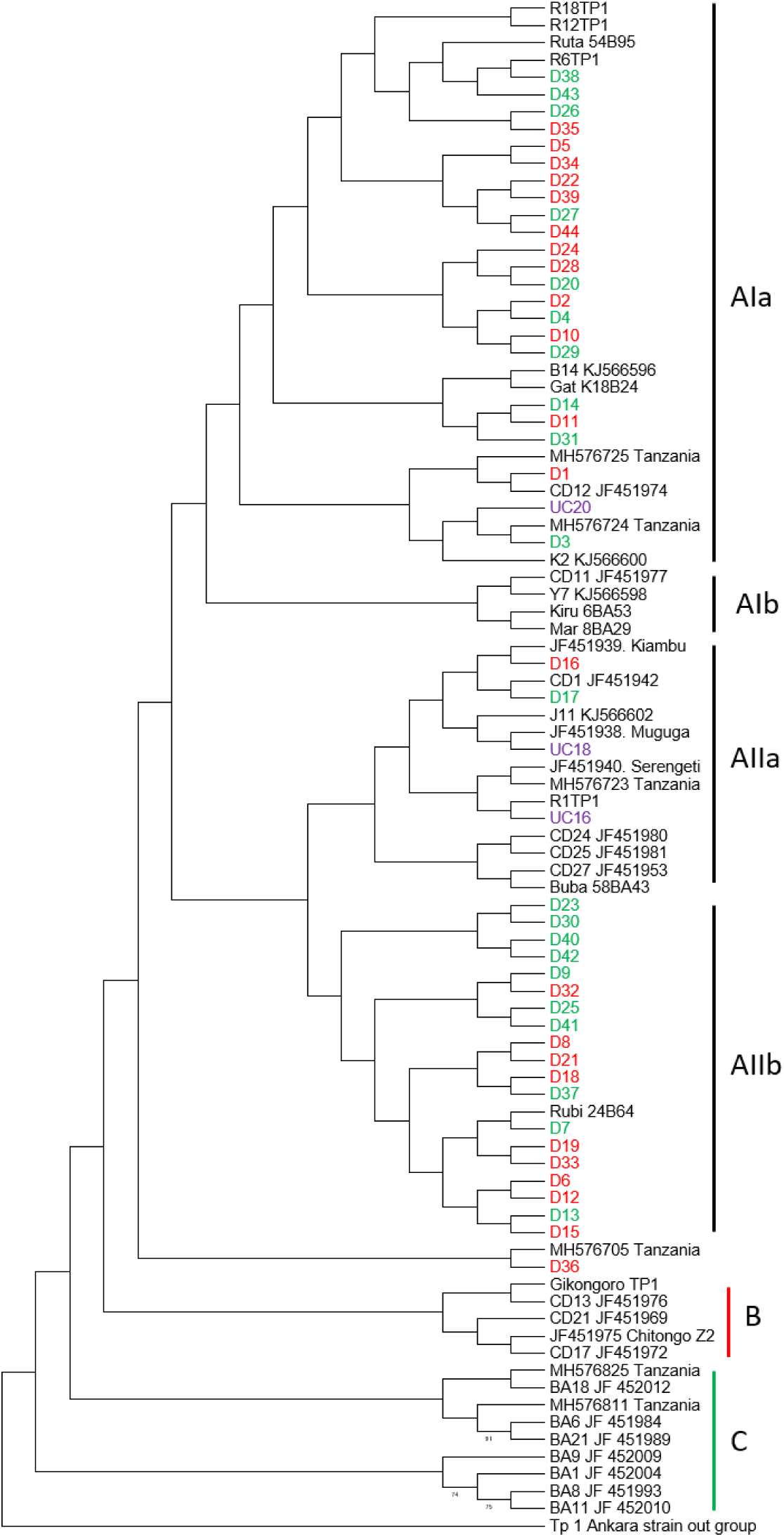
*T. parva* Tp1 phylogenetic tree. The tree was constructed from 432 bp nucleotide sequences using MEGA version 6 with 1000 bootstrap replicates for confidence. Samples from sentinel, immunised and control groups are bold and purple, green and red, respectively.

Analysis of Tp1 haplotypes using network 10 showed a star-like pattern with nine haplotypes (H11, H16, H15, H22, H18, H8, H23, H9, H8) radiating from haplotype H7 (Fig. 4), indicating the possibility of population expansion. H7, mainly comprising samples from the control group, was the most abundant haplotype (n=15). H1, despite being one of the minor haplotypes, included sequences from the sentinel, control, and immunized groups. All other haplotypes either comprised one group (e.g. H22 with control group samples) or included samples from both the control and immunized groups. The MC haplotypes were least represented overall, suggesting that the majority of samples from both the control and immunized groups differed from the MC (Fig. 4).

**Fig. 4:**
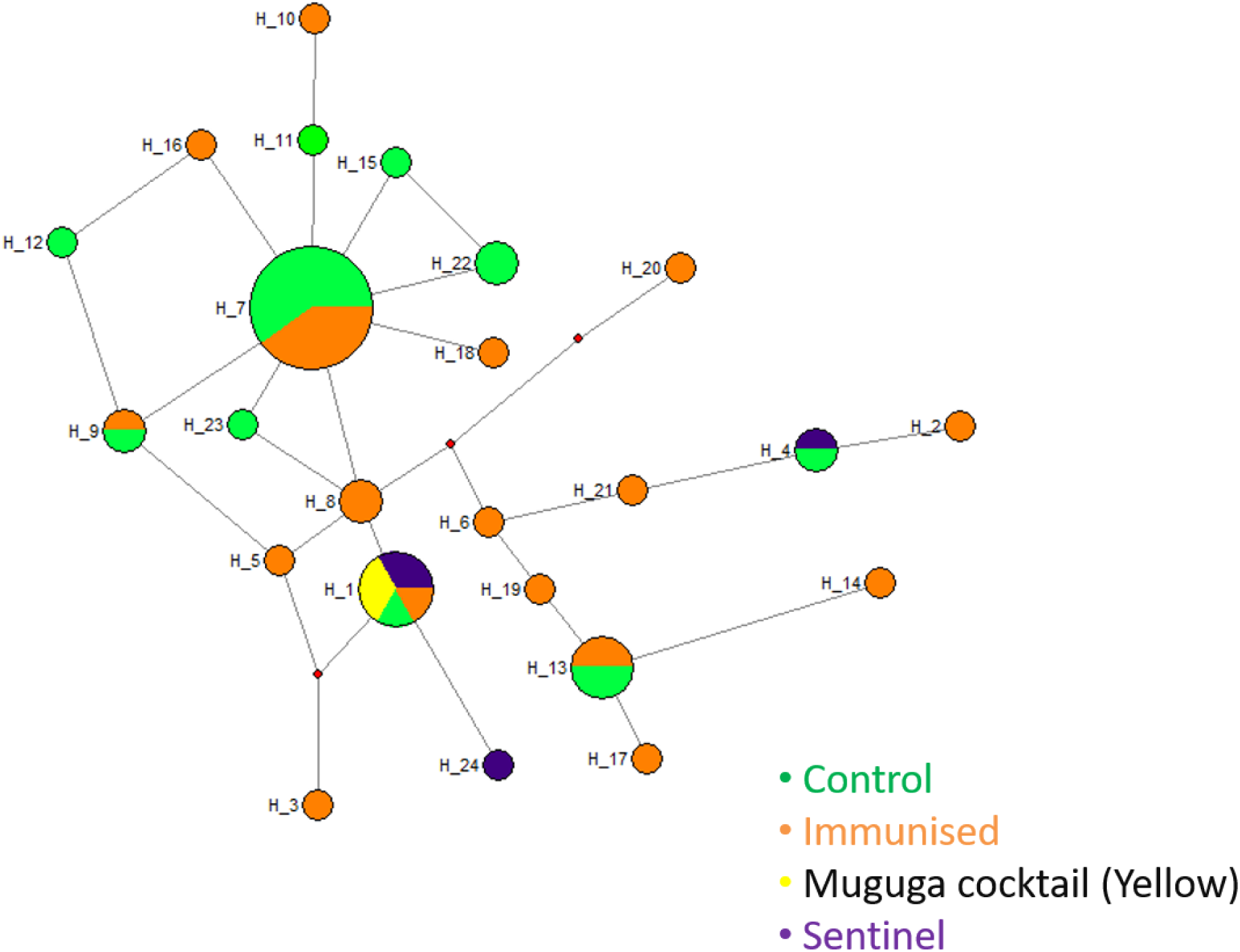
Median-joining network of the *T.parva* Tp1 gene, constructed using Network 5 and based on the polymorphic sites of Tp1. The network displays a star-like radiation pattern, with circle sizes corresponding to haplotype frequency and color codes indicating the origin of the samples.

### Tp2 Locus

The Tp2 gene encodes a 174-amino acid protein that consists of six epitopes: CTL 1 (SHEELKKLGML), CTL 2 (DGFDRDALF), CTL 3 (KSSHGMGKVGK), CTL 4 (FAQSLVCVL), CTL 5 (QSLVCVLMK), and CTL 6 (KTSIPNPCKW). The complete Tp2 gene (531 bp) of *T. parva* was sequenced from samples in the sentinel, immunized, and control groups. Regarding CTL 1, no MC epitopes were identified in both control and immunized samples; instead, ten (10) variants were identified (Table 7). MC epitopes were only identified on CTL 4, 5, and 6 in 6, 6, and 11 post-immunization samples, respectively. The observed MC variants were 10, 10, 7, 9, 10, and 7 on epitopes 1 to 6, respectively. Out of a total of 258 epitopes identified across all loci, only 23 were similar to MC epitopes, while the majority (n=235) were variants of the MC epitope (Table 7). Additionally, all epitopes identified from sentinel sequences were variants of the MC epitope. Overall, the MC epitope was poorly represented across all six epitopes of the Tp2 protein (Table 7). A DNA polymorphism score of 13.2%, indicating high nucleotide polymorphism, was calculated from the sentinel, control, and immunized samples. The mean ratio of dN/dS was 0.659 with 15 purifying selection sites.

**Table 7:**
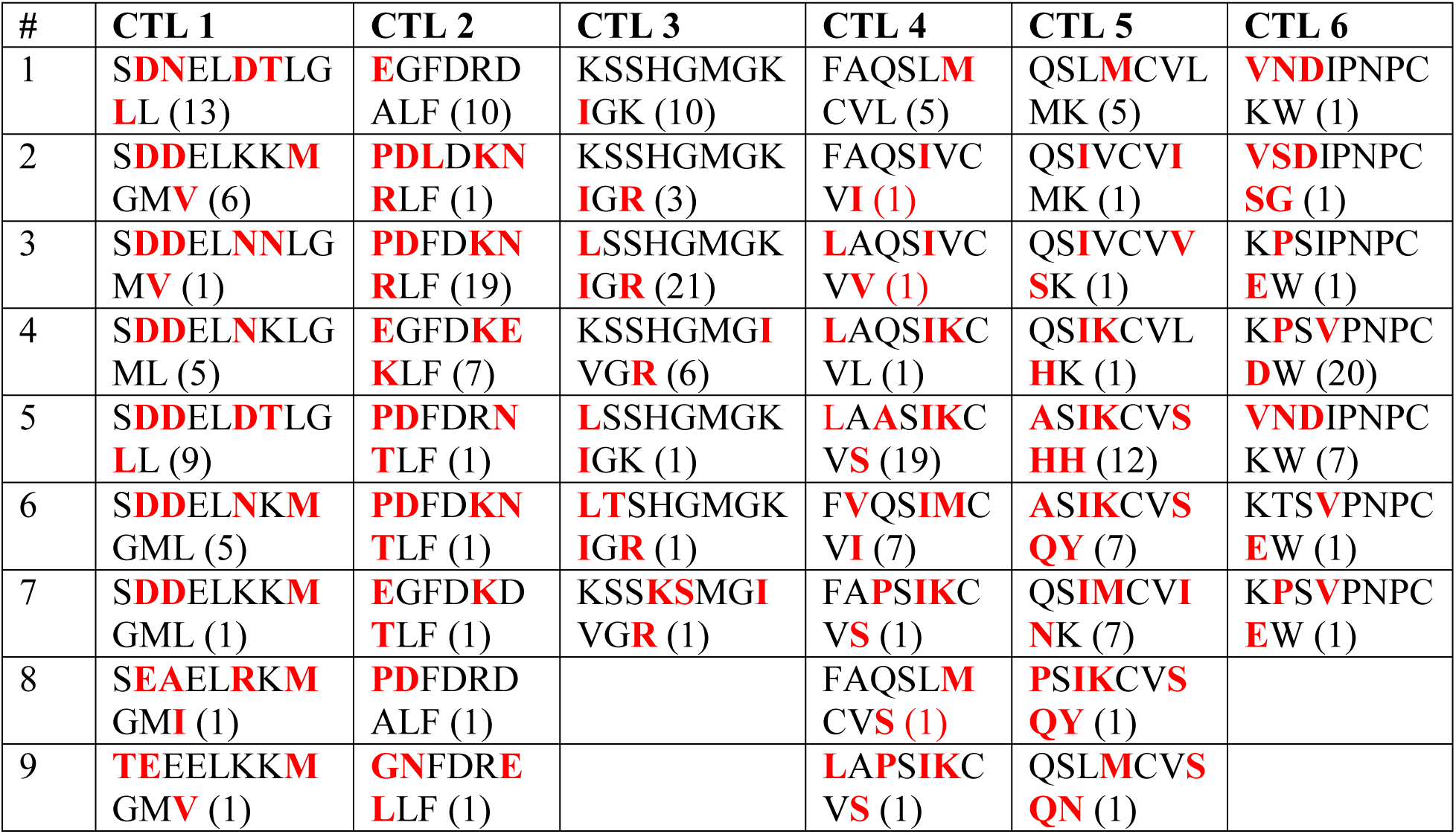
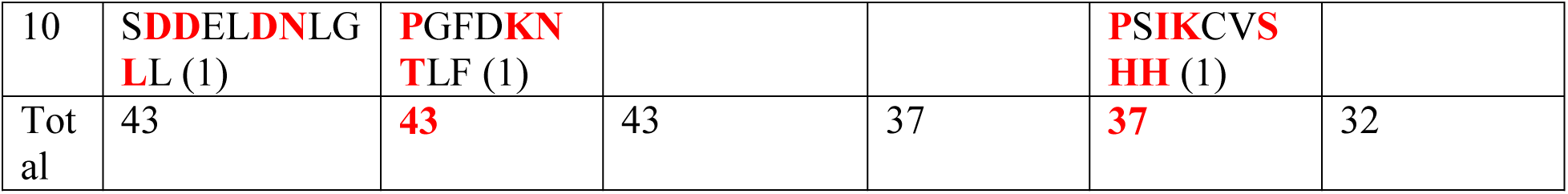
Tp2 epitope variants identified in samples from eastern DRC.

Furthermore, the phylogenetic analysis of the Tp2 gene (Fig. 5) unveiled three primary clusters, with sequences from this study grouping within clusters D and E. Cluster D further divided into two sub-clusters: DI, associated with the Muguga cocktail strains, and DII, linked with the chitongo strain (Fig. 5). Cluster DI encompassed 11 study sequences (6 immunized and 5 control), while cluster DII and cluster E comprised 23 (12 immunized, 10 controls, and 1 sentinel) and 9 (5 immunized, 2 controls, and 2 sentinels) sequences, respectively. In summary, the phylogenetic analysis revealed that the majority of the study sequences did not closely cluster with the Muguga cocktail strains (Fig. 5).

**Fig. 5:**
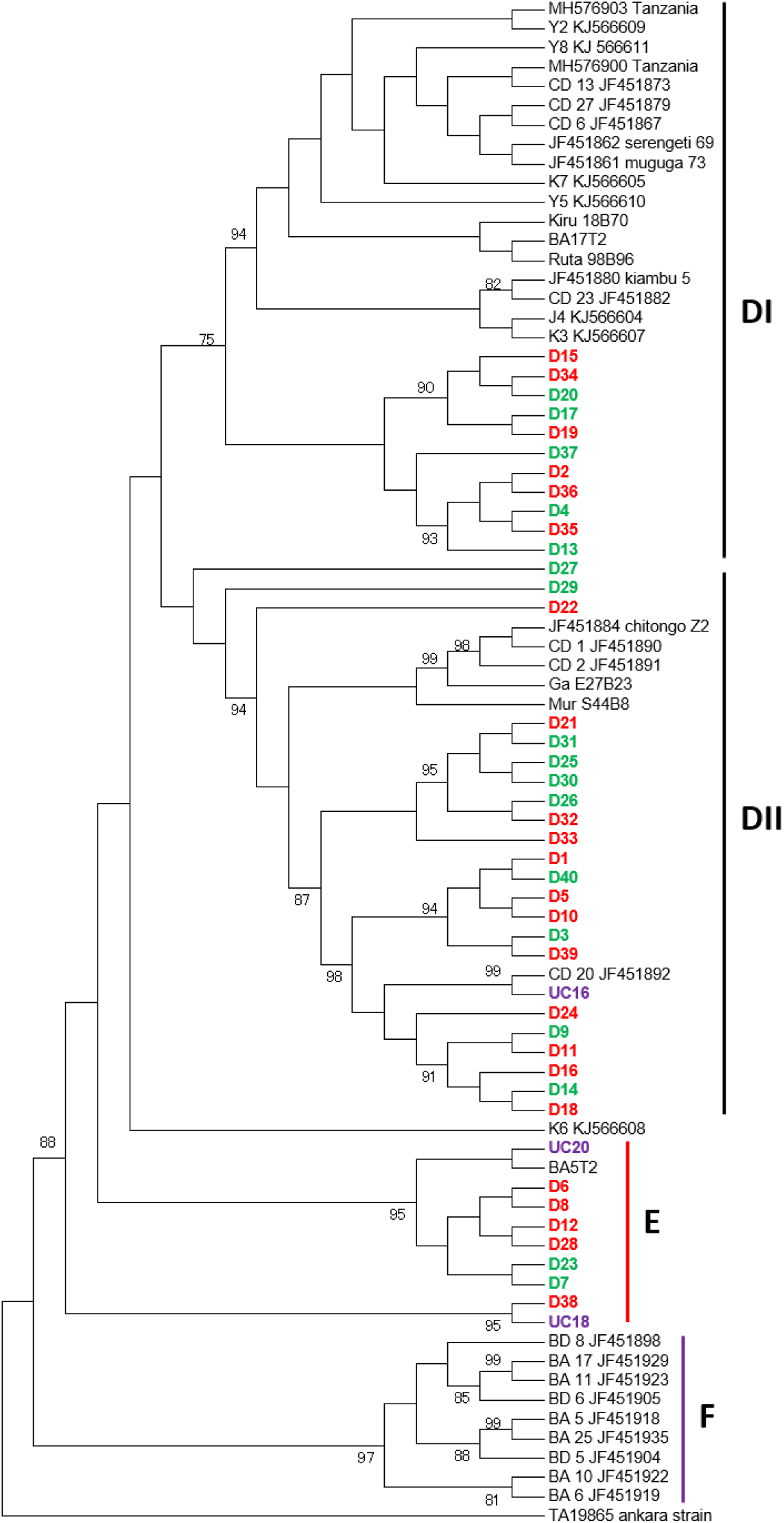
*T. parva* Tp1 phylogenetic tree. The tree was constructed from 531 bp nucleotide sequences using MEGA version 6 with 1000 bootstrap replicates as a measure of confidence. Sentinel, immunised and control sequences are given in bold and purple, green and red, respectively.

The analysis of haplotype diversity revealed distinctive and non-shared haplotypes in the MC and sentinel samples, which were not shared among the immunized or control groups (Fig. 6). Specifically, Muguga cocktail haplotypes H1 and H2 exhibited a direct relation only to H8, encompassing both immunized and control group samples. Within the sentinel haplotypes, only haplotype H17 showed a direct connection to H13, involving both immunized and control group sequences. The remaining sentinel haplotypes were indirectly linked to other haplotypes, such as H15, through median vectors. In summary, network analysis demonstrated that sentinel haplotypes were not connected to the MC haplotypes, nor were they associated with the haplotypes found in both the immunized and control groups.

**Fig. 6:**
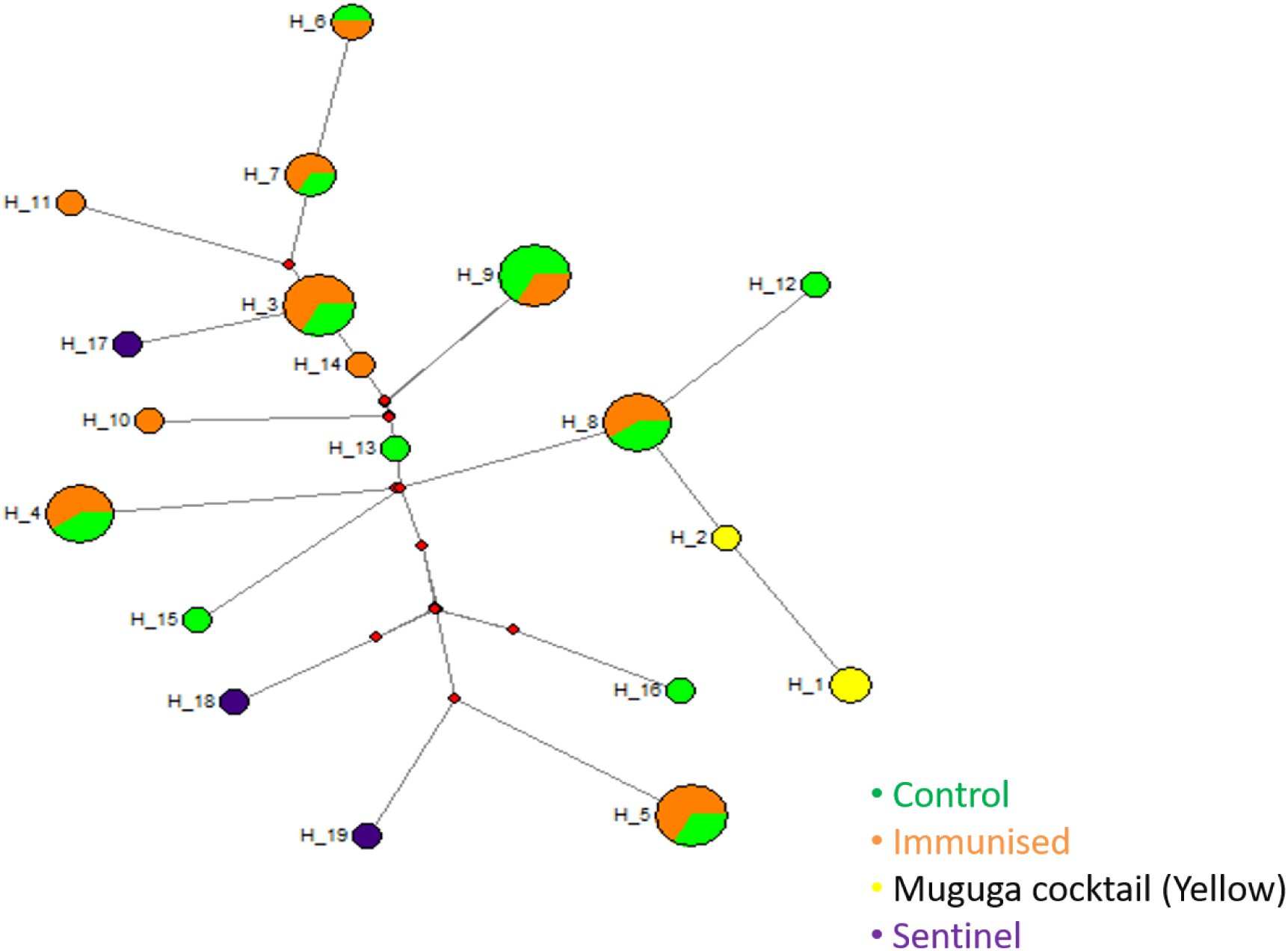
Median-joining network of the *T.parva* Tp2 gene constructed using Network 5 and based on the polymorphic sites of Tp2. The size of the circle corresponds to haplotype frequency and the color codes indicate the origin of the samples.

### Marker diversity and allelic variation

Samples from control, immunized, and sentinel cattle from the DRC were genotyped using a panel of five microsatellites (MS39, MS25, MS7, MS33, and MS19) and one mini-satellite marker (ms9), covering the four chromosomes of *T. parva*. Among the markers, MS39 was the most polymorphic, producing 25 alleles, while MS19 was the least polymorphic, identifying only 13 alleles (Table 8). High genetic diversities were also observed in sentinel, control and immunized populations (Table 8).

**Table 8:**
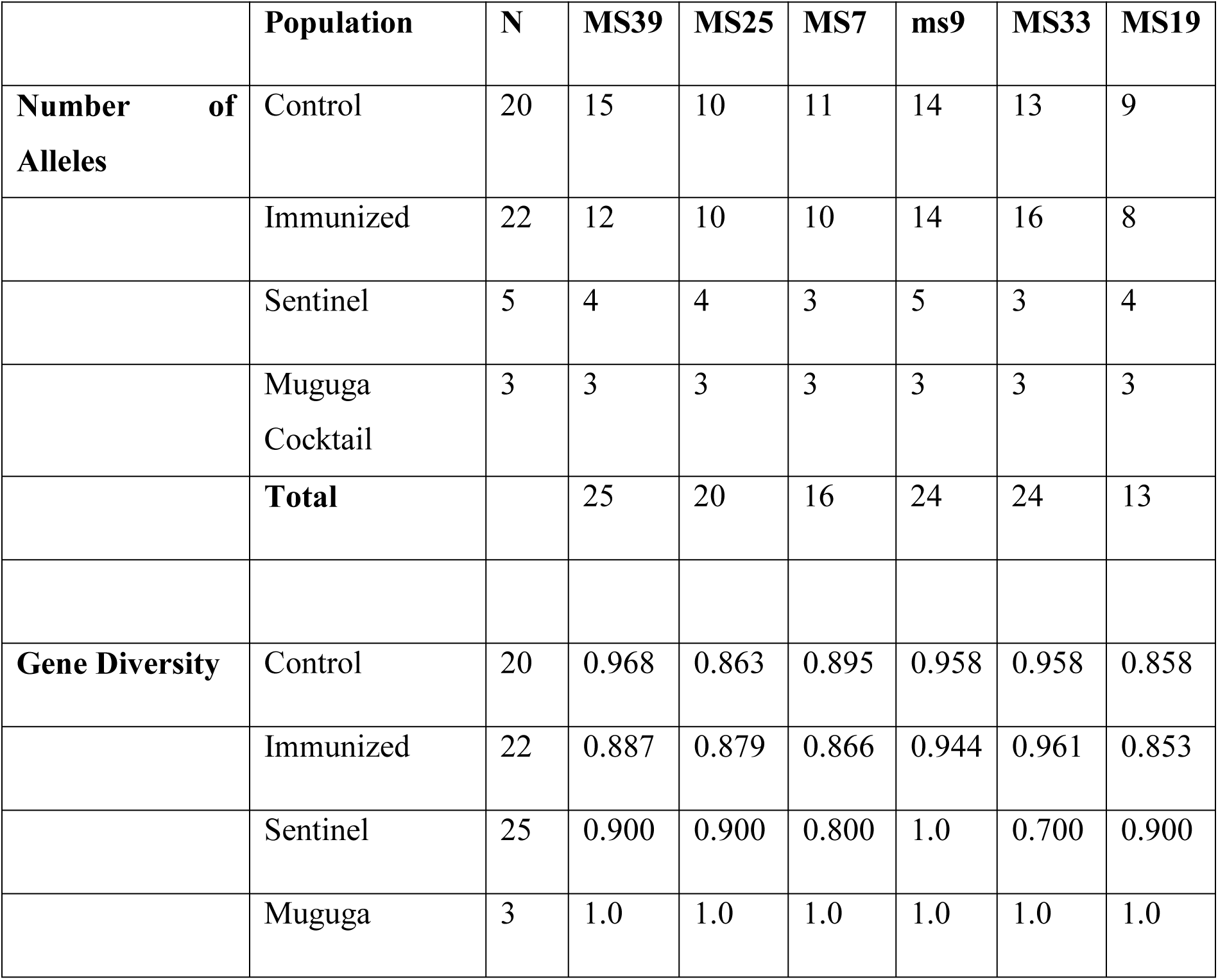
Alleles identified and genetic diversity in sentinel, immunized and control groups.

Across the six loci, there were both shared and unique alleles. The MC exhibited a total of 18 alleles, with 14 being unique to MC and 4 shared with the control, immunized, and sentinel populations. The control and immunized populations each possessed a total of 35 and 30 unique alleles, respectively. When considering all populations together, the total number of unique alleles across the six loci was 91, while the number of shared alleles was 38. The greater proportion of unique alleles indicated genetic sub-structuring among the populations (Fig. 7).

**Fig. 7.**
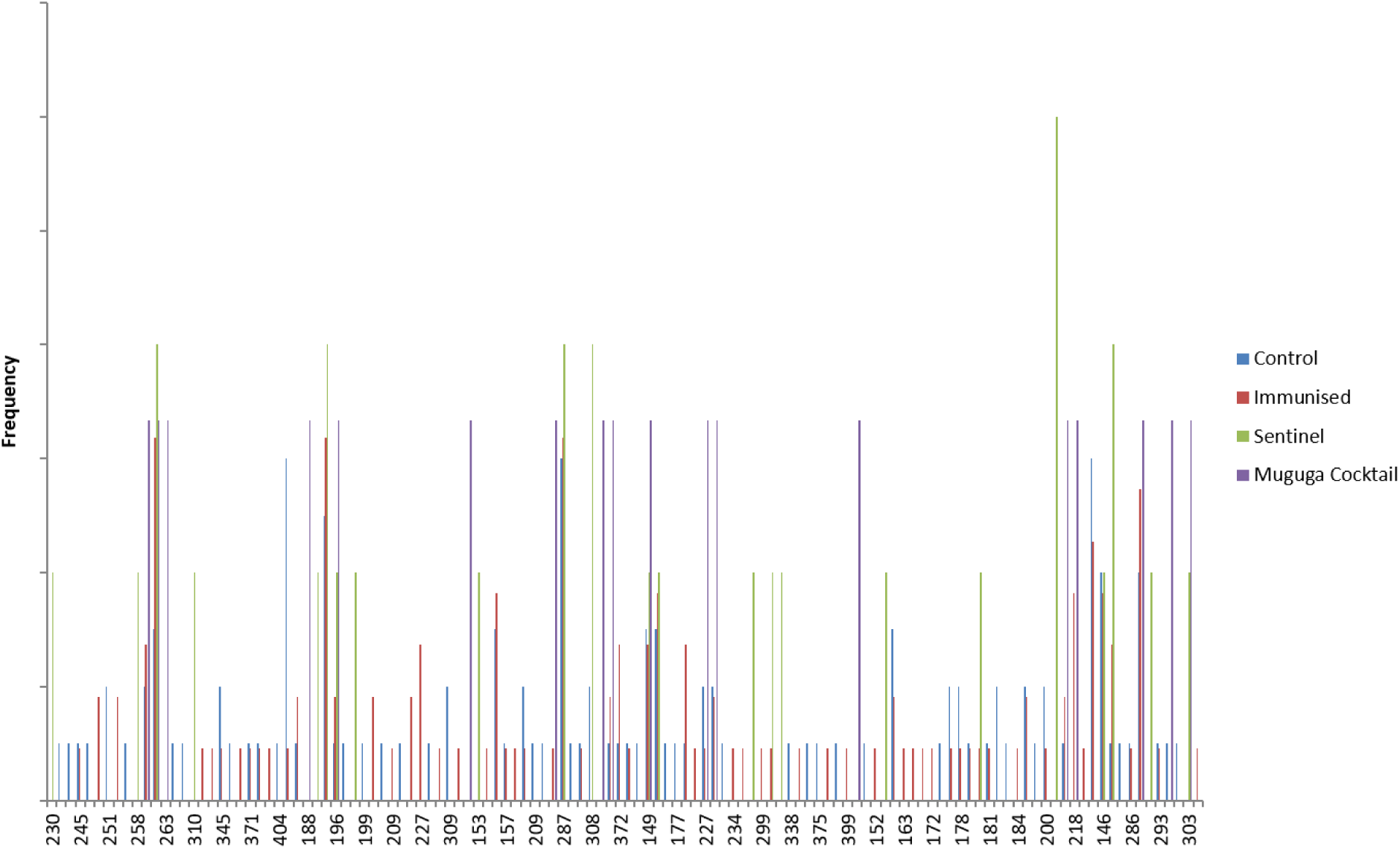
Overall frequency of alleles from sentinel, immunised, control and vaccine groups. The dataset presents shared and unique alleles determined by calculating the proportions of the predominant alleles relative to the total of each satellite marker. Histograms were generated using the multi-locus genotype data.

### Similarity analysis, population differentiation and diversity

To evaluate the observed genetic sub-structuring depicted in Fig. 7, Principal Component Analysis (PCA) was employed. The PCA results revealed that samples from the Democratic Republic of the Congo (DRC) occupied all four quadrants, whereas the Muguga cocktail (MC) samples were confined to a single quadrant (Fig. 8). Notably, thirty-six control (n=13), immunized (n=18) and sentinel (n=5) samples closely clustered with the MC in cluster A, while the remaining eleven (11) were divided between cluster B (n=6) and C (n=5), comprising of both control and immunized samples. This cluster pattern indicates a degree of sub-structuring. The degree of sub-structuring, as illustrated in Fig. 7 and Fig. 8, was further assessed using Wright’s F index. Fst values of 0.119 and 0.096 were obtained when cluster population A, (MC, control, sentinel and immunized samples), was compared to cluster populations B and C, respectively. When all populations were treated as one, an Fst value of 0.118 was obtained.

**Fig. 8:**
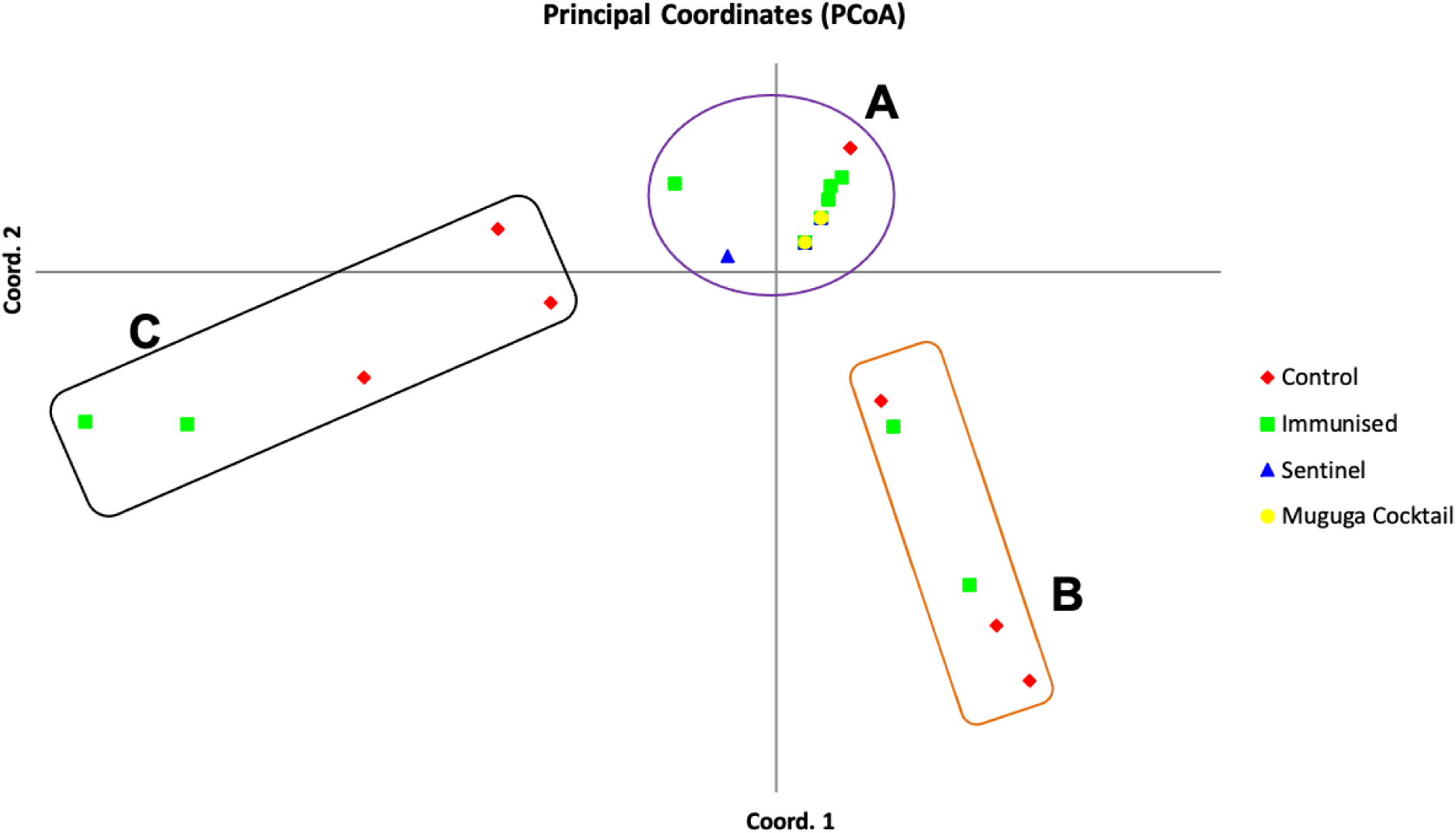
Principal Component Analysis (PCA) for sentinel, immunised, control and muguga cocktail vaccine samples revealing sub-structuring among three populations: Population A includes muguga cocktail, immunised, control and sentinel samples; Population B and C comprises of immunised and control samples. The numbers on each axis indicate the proportion of variance explained by the population dataset.

## Discussion

Molecular techniques, such as PCR and DNA sequencing, have revolutionized the detection of previously unidentified pathogens in regions believed to be pathogen-free. To establish successful immunization programs and ensure optimal protection of the local cattle population, it is imperative to identify suitably matched vaccine strains of *T. parva* for use in new areas. Molecular-based studies on prevailing *T. parva* populations are essential for this purpose, as the infection and treatment method only confer strain-specific immunity (34,35). Positive outcomes from similar studies in Rwanda (25) and Burundi (13) underscore the potential success of this approach. To assess the distribution of *T.parva* and its vector, *R.appendiculatus*, and to evaluate similarities between *T.parva* populations in Kabasha village, eastern DRC, and the Muguga cocktail vaccine stocks, this study employed a comprehensive approach that included a knowledge and perception survey among farmers, alongside analyses of prevalence, tick distribution, sequences, and microsatellites. The study also included a rigorous immunization and field challenge trial aimed at informing and developing effective immunization strategies against East Coast fever (ECF) in the eastern DRC region. Notably, this is the first study to utilize the sequence diversity of *T.parva* CTL antigens Tp1 and Tp2, along with mini- and microsatellite analysis and field challenge trials, to characterize and compare field samples with known vaccine isolates in the DRC.

This study strategically focused on three key variables; age group, sex, and agro-ecological zones while sampling a total of 756 cattle across three provinces. The majority of sampled animals were adult females, aligning with common slaughtering practices involving male calves. The study revealed the presence of five tick species (Table 2), with *R. appendiculatus* standing out as the most prevalent at 68.04%, signifying a potential risk of Theileriosis transmission in the region. Tick distribution varied among provinces, with South-Kivu exhibiting the highest tick burden per animal. Notably, the high prevalence of *R. appendiculatus* correlated with a 27.5% occurrence of *T. parva*, as determined through ELISA and PCR methods (Tables 3 and 4).

To identify the strains of prevailing *T. parva* populations and evaluate the effectiveness of the available Muguga cocktail vaccine, five sentinel cattle were stationed in Kabasha village for 20 days. During this period, four out of the five animals developed a severe form of East Coast fever (ECF), leading to the selection of Kabasha village as the site for subsequent immunization and challenge trials. This observation highlighted the critical importance of Kabasha village in understanding and addressing the dynamics of *T. parva* strains in the region. Subsequently, a field challenge trial was conducted in Kabasha village involving 100 naïve cattle. These animals were vaccinated with the available Muguga cocktail vaccine and allowed to graze freely in the village. After the exposure period, all the control animals and more than half of the vaccinated animals succumbed to ECF, resulting in an immunization success rate of only 46.5%. This indicated a failure of the vaccine to provide adequate protection against ECF and likely that the available stocks may not offer cross protection again the Kabasha village strain of *T. parva*.

To further understand the characteristics of the *T. parva* population in Kabasha village and how it compares with the Muguga cocktail stocks, molecular analyses were conducted. Phylogenetic analysis of the p67 gene revealed the exclusive presence of cattle-derived *T. parva* strains, all possessing allele type 1, as observed in other studies (13). Analysis of the CTL Tp1 (Fig. 3) and Tp2 (Fig. 5) antigens showed three clusters, contrasting with findings from other studies (12–15,25). The sentinel samples (2/3) were closely related to the Muguga cocktail based on Tp1 and exhibited distance relatedness to Muguga cocktail on Tp2 (Fig. 5). On both Tp1 and Tp2 phylogenetic analyses, immunized and control samples separated into two clusters. Within these clusters, both immunized and control samples were present, implying the presence of similar strains in both groups. On Tp2, two sentinel and control samples and five immunized samples formed a separate cluster, suggesting they could have originated from a different population. The majority of samples from the sentinel, immunized, and control groups didn’t share the same antigenic profile with the Muguga cocktail, especially considering Tp2 epitopes (Table 7). This was evident from the high proportion of epitope variants on Tp2 (Table 7), despite the majority of epitopes on Tp1 being similar to Muguga cocktail. This discrepancy may be attributed to differences in the conservation nature of both genes.

Ultimately, the implication is that most *T. parva* field populations, while appearing Muguga cocktail-like, might not share the same immunological response and could constitute a different population from the Muguga cocktail. This is further supported by the few haplotypes that were shared between the sentinel, immunized, and control populations and the Muguga cocktail vaccine stocks (Fig. 4 and 6). This observation aligns with previous studies in different regions (13,25,36). Additionally, evidence of expanding populations with similar haplotypes across the region suggests the possibility of open grazing, coupled with free trade in animals or movement among local farmers, driving the spread of *T. parva* infection. This practice hampers the implementation of adequate control strategies, such as frequent dipping and livestock movement bans, crucial in mitigating the effects of theileriosis.

It is important to note that at the time of sampling, part of this region was experiencing armed conflict; therefore, the samples used for molecular analysis were obtained from the same place (Kabasha village). Samples that appear different from the Muguga cocktail may represent breakaway populations that do not succumb to ITM using Muguga cocktail. Nevertheless, sequence analysis alone is insufficient to determine the most appropriate vaccine strain for field challenge trials with respect to *T. parva*. Due to the highly conserved nature of Tp1 and diversity of Tp2 genes, determining the similarity of proposed vaccine isolates to field populations is challenging. To enhance result resolution, population genetic analysis involving six (6) sites on the *T. parva* genome was conducted. Population genetic analysis revealed high gene diversities in all populations across all loci (Table 8), indicating the presence of a diverse population. Despite the lack of close clustering with MC vaccine isolates on phylogenetic and network analysis, sentinel, immunized, and control samples also did not share the majority of alleles on microsatellite analysis (Fig. 7). In contrast, PCA (Fig. 8) showed close clustering of samples comprising sentinel, immunized, and control populations, with a few clustering independently. This could be attributed to detecting Muguga cocktail vaccine strains, along with control samples infected with Muguga cocktail-like parasites. Furthermore, based on sequence diversity and population genetic data, it is likely that more than one population similar to the Muguga cocktail but genetically sub-structured from it is present at the study site. This is evidenced by the breakaway populations observed in clusters B and C on the PCA (Fig. 8) and the Fst values of 0.119 and 0.096 for comparisons of A vs. B and A vs. C, respectively. An Fst value of 0.118 when all populations are treated as a single population indicates moderate genetic differentiation among the populations.

In conclusion, this study revealed the prevalence of *Rhipicephalus appendiculatus* ticks, a potential vector for Theileriosis transmission, with tick distribution varying among DRC provinces. While some populations appeared Muguga cocktail-like, significant differences in antigenic profiles were observed, indicating the presence of distinct populations. Despite the armed conflict in the region during sampling, the study provided valuable insights into the genetic diversity of *T. parva* populations, suggesting the presence of diverse populations. The immunization and field challenge trial indicated a low survival rate, underscoring the limitations of the current vaccine against field strains in eastern DRC. The study concludes that more than one population similar to the Muguga cocktail may be present in the study site, as observed in clusters B and C on PCA. Nevertheless, this study contributes significantly to the understanding of *T. parva* dynamics in the region, emphasizing the complexities of vaccine strain selection and the importance of continuous monitoring and adaptation of control strategies in the face of evolving parasite populations.

### Ethical approval and Consent to participate

Ethical clearance for the recombinant DNA experiments conducted in this study received approval from the University of Zambia Biomedical Research Ethics Committee (UNZABREC) under REF. 233-2019. Permission for sample collection and field trial was obtained from the Department of Agriculture, Livestock, and Fisheries (AGRIPEL) in the DRC. It is essential to note that the animals underwent humane handling procedures throughout the sample collection and the immunization/field challenge study.

### Consent for publication

Not applicable.

## Availability of data and materials

All the nucleotide sequences obtained and used in this study have been deposited in the DNA database of Japan with accession numbers LC863893-LC863940. All other material is contained within the manuscript.

## Competing interests

WM and DKA contributed equally to this work while other authors declare no competing interests.

## Funding

This publication has been supported by the Global Alliance for Livestock and Veterinary medicines (GALV*med*) with funding from the Bill & Melinda Gates Foundation and UK aid from the UK Government. The findings and conclusions contained within this publication are those of the authors and do not necessarily reflect the positions or policies of the Bill & Melinda Gates Foundation nor the UK Government. Financial support for this work was within the framework of the project on the characterization and population genetics of *Theileria parva* strains in eastern, central and southern Africa (UOZ-R34A0548A3).

## Author contributions

Experimental design, data analysis, writing manuscript draft, W.M. and D.K.A.; field sample collection, V.M., K.M.K., K.M.H.; tick identification, K.M.K.; immunisation and challenge study, A.J.M., D.K.A. and W.M.; laboratory work, S.M., W.M. and D.K.A.; drafting and editing manuscript, V.M., S.M. and J.S.; Experimental design and manuscript draft editing, B.N., and A.J.M. All authors have read and approved the final version of the manuscript.

## Acknowledgements

We thank the veterinary assistants and farmers from the three Provinces of North Kivu, South Kivu and Ituri for their role in sample collection. We also thank the technical staff at the Centre for Ticks and Tick-Borne Diseases, Lilongwe, Malawi and University of Zambia, School of Veterinary Medicine for their role in some of the laboratory work. Furthermore, we are grateful to Phil Toye and Roger Pelle of the International Livestock Research Institute, Nairobi, Kenya for their helpful advice at the conception of the idea phase.

## Additional File 1. Satellite markers used to genotype field samples from Eastern DR Congo

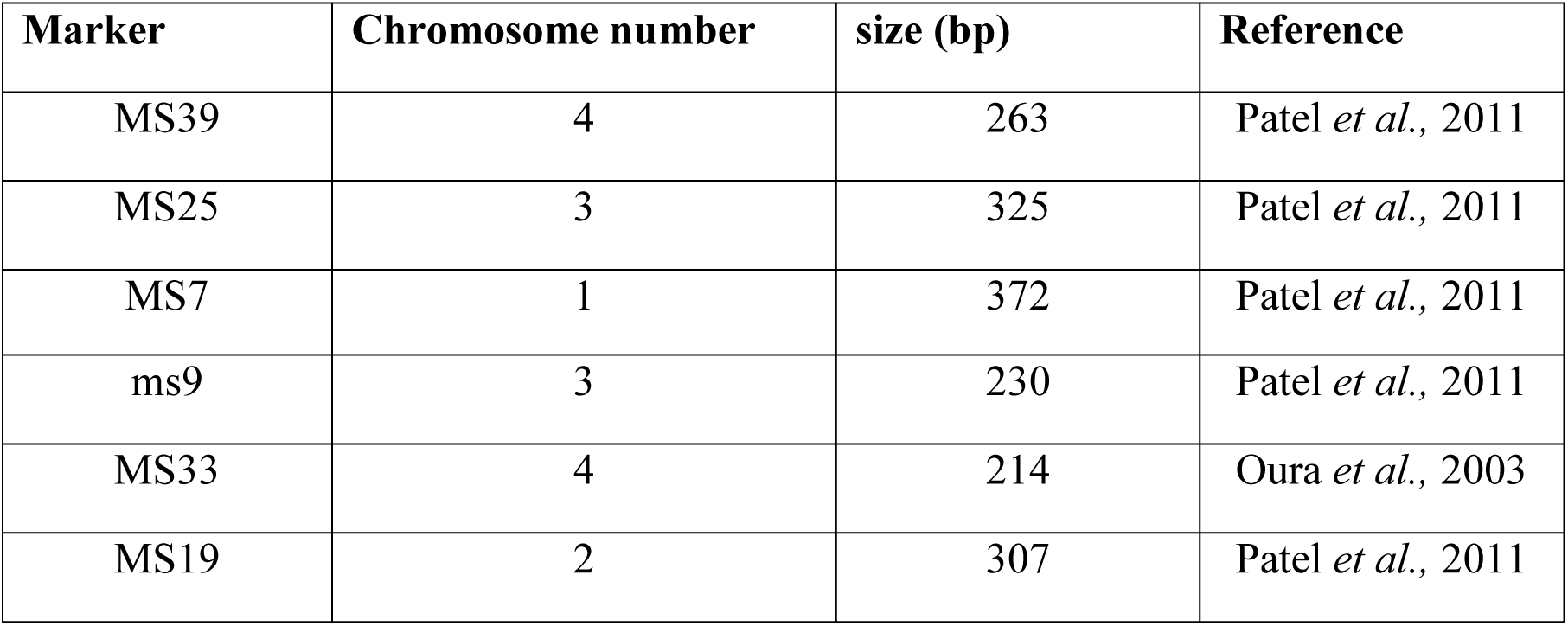

| **Marker** | **Chromosome number** | **size (bp)** | **Reference** |
| --- | --- | --- | --- |
| MS39 | 4 | 263 | Patel *et al.*, 2011 |
| MS25 | 3 | 325 | Patel *et al.*, 2011 |
| MS7 | 1 | 372 | Patel *et al.*, 2011 |
| ms9 | 3 | 230 | Patel *et al.*, 2011 |
| MS33 | 4 | 214 | Oura *et al.*, 2003 |
| MS19 | 2 | 307 | Patel *et al.*, 2011 |

## Notes

### Competing Interest Statement

Walter Muleya and David Kalenzi Atuhaire contributed equally to this work

## References

1. Lessard P, L’Eplattenier R, Norval RA, Perry BD, Dolan TT, Burrill A, et al. The use of geographical information systems in estimating East Coast fever risk to African livestock. Acta Vet Scand Suppl [Internet]. 1988;84:234–6. Available from: http://www.ncbi.nlm.nih.gov/pubmed/3232616

2. Muraguri GR, Kiara HK, McHardy N. Treatment of East Coast fever: a comparison of parvaquone and buparvaquone. Vet Parasitol. 1999;87(1):25–37.

3. Schein E, Warnecke M, Kirmse P. Development of Theileria parva (Theiler, 1904) in the gut of Rhipicephalus appendiculatus (Neumann, 1901). Parasitology. 1977;75(3):309–16.

4. Lessard P, L’Eplattenier R, Norval RA, Kundert K, Dolan TT, Croze H, et al. Geographical information systems for studying the epidemiology of cattle diseases caused by Theileria parva. Vet Rec [Internet]. 1990;126(11):255–62. Available from: http://www.ncbi.nlm.nih.gov/pubmed/2327044

5. Makumyaviri AM, Mwilambwe KP. Dépistage et diagnostic dela theilériose et del’anaplasmose chez les bovins élevés au ranchdes Marungu, République Démocratique du Congo. Cah Vét Congo. 1988;01:22–23.

6. Kalume MK, Saegerman C, Mbahikyavolo DK, Makumyaviri AM, Marcotty T, Madder M, et al. Identification of hard ticks (Acari: Ixodidae) and seroprevalence to Theileria parva in cattle raised in North Kivu Province, Democratic Republic of Congo. Parasitol Res. 2013 Feb;112(2):789–97.

7. Mehlhorn H, Shein E. The piroplasms: life cycle and sexual stages. Adv Parasitol. 1984;23:37–103.

8. Pegram RG, Lemche J, Chizyuka HGB, Sutherst RW, Floyd RB, Kerr JD, et al. Ecological aspects of cattle tick control in central Zambia. Med Vet Entomol. 1989;3(3):307–12.

9. Radley DE. Infection and Treatment Method of Immunization Against Theileriosis. In: Advances in the Control of Theileriosis. Springer Netherlands; 1981. p. 227–37.

10. Radley DE, Brown CGD, Burridge MJ, Cunningham MP, Kirimi IM, Purnell RE, et al. East coast fever: 1. Chemoprophylactic immunization of cattle against Theileria parva (Muguga) and five theilerial strains. Vet Parasitol. 1975;1(1):35–41.

11. Oura CAL, Bishop R, Asiimwe BB, Spooner P, Lubega GW, Tait A. Theileria parva live vaccination: parasite transmission, persistence and heterologous challenge in the field. Parasitology. 2007;134(9):1205–13.

12. Chatanga E, Kainga H, Maganga E, Hayashida K, Katakura K, Sugimoto C, et al. Molecular identification and genetic characterization of tick-borne pathogens in sheep and goats at two farms in the central and southern regions of Malawi. Ticks Tick Borne Dis. 2021 Mar;12(2):101629.

13. Atuhaire DK, Muleya W, Mbao V, Niyongabo J, Nyabongo L, Nsanganiyumwami D, et al. Molecular characterization and population genetics of Theileria parva in Burundi’s unvaccinated cattle: Towards the introduction of East Coast fever vaccine. Rosenthal BM, editor. PLoS One. 2021 May;16(5):e0251500.

14. MacHugh ND, Connelley T, Graham SP, Pelle R, Formisano P, Taracha EL, et al. CD8 + T-cell responses to Theileria parva are preferentially directed to a single dominant antigen: Implications for parasite strain-specific immunity. Eur J Immunol. 2009 Sep;39(9):2459–69.

15. Pelle R, Graham SP, Njahira MN, Osaso J, Saya RM, Odongo DO, et al. Two Theileria parva CD8 T Cell Antigen Genes Are More Variable in Buffalo than Cattle Parasites, but Differ in Pattern of Sequence Diversity. Langsley G, editor. PLoS One. 2011 Apr;6(4):e19015.

16. Oura CA, Odongo D, Lubega G, Spooner P, Tait A, Bishop R. A panel of microsatellite and minisatellite markers for the characterisation of field isolates of Theileria parva. Int J Parasitol. 2003;33(14):1641–53.

17. Oura CAL, Asiimwe BB, Weir W, Lubega GW, Tait A. Population genetic analysis and sub-structuring of Theileria parva in Uganda. Mol Biochem Parasitol. 2005;140(2):229–39.

18. Odongo D, Oura C, Spooner P, Kiara H, Mburu D, Hanotte O, et al. Linkage disequilibrium between alleles at highly polymorphic mini- and micro-satellite loci of Theileria parva isolated from cattle in three regions of Kenya. Int J Parasitol. 2006;36(8):937–46.

19. Muleya W, Namangala B, Simuunza M, Nakao R, Inoue N, Kimura T, et al. Population genetic analysis and sub-structuring of Theileria parva in the northern and eastern parts of Zambia. Parasit Vectors. 2012 Dec;5(1):255.

20. Asiimwe BB, Weir W, Tait A, Lubega GW, Oura CAL. Haemoparasite infection kinetics and the population structure of Theileria parva on a single farm in Uganda. Vet Parasitol. 2013;193(1–3):8–14.

21. Geysen D, Bazarusanga T, Brandt J, Dolan TT. An unusual mosaic structure of the PIM gene of Theileria parva and its relationship to allelic diversity. Mol Biochem Parasitol. 2004 Feb;133(2):163–73.

22. Katzer F, Ngugi D, Walker AR, McKeever DJ. Genotypic diversity, a survival strategy for the apicomplexan parasite Theileria parva. Vet Parasitol. 2010;167(2–4):236–43.

23. Nene V. Organisation and informational content of the Theileria parva genome. Mol Biochem Parasitol. 1998;95(1):1–8.

24. Katende J, Morzaria S, Toye P, Skilton R, Nene V, Nkonge C, et al. An enzyme-linked immunosorbent assay for detection of Theileria parva antibodies in cattle using a recombinant polymorphic immunodominant molecule. Parasitol Res. 1998;84(5):408–16.

25. Atuhaire DK, Muleya W, Mbao V, Bazarusanga T, Gafarasi I, Salt J, et al. Sequence diversity of cytotoxic T cell antigens and satellite marker analysis of Theileria parva informs the immunization against East Coast fever in Rwanda. Parasit Vectors. 2020;13(1):452.

26. Pelle R, Graham SP, Njahira MN, Osaso J, Saya RM, Odongo DO, et al. Two Theileria parva CD8 T Cell Antigen Genes Are More Variable in Buffalo than Cattle Parasites, but Differ in Pattern of Sequence Diversity. PLoS One. 2011;6(4):e19015.

27. Tamura K, Stecher G, Peterson D, Filipski A, Kumar S. MEGA6: Molecular Evolutionary Genetics Analysis Version 6.0. Mol Biol Evol. 2013 Dec;30(12):2725–9.

28. Librado P, Rozas J. DnaSP v5: a software for comprehensive analysis of DNA polymorphism data. Bioinformatics. 2009;25(11):1451–2.

29. Pond SLK, Poon AFY, Frost SDW. Estimating selection pressures on alignments of coding sequences. In: The Phylogenetic Handbook. Cambridge University Press; 2009. p. 419–90.

30. Delport W, Scheffler K, Gravenor MB, Muse S V., Kosakovsky Pond S. Benchmarking Multi-Rate Codon Models. Poon AFY, editor. PLoS One. 2010 Jul;5(7):e11587.

31. Peakall R, Smouse PE. genalex 6: genetic analysis in Excel. Population genetic software for teaching and research. Mol Ecol Notes. 2006 Mar;6(1):288–95.

32. Nene V, Musoke A, Gobright E, Morzaria S. Conservation of the sporozoite p67 vaccine antigen in cattle-derived Theileria parva stocks with different cross-immunity profiles. Infect Immun. 1996;64(6):2056–61.

33. Sibeko KP, Geysen D, Oosthuizen MC, Matthee CA, Troskie M, Potgieter FT, et al. Four p67 alleles identified in South African Theileria parva field samples. Vet Parasitol. 2010 Feb;167(2–4):244–54.

34. Taracha EL, Goddeeris BM, Morzaria SP, Morrison WI. Parasite strain specificity of precursor cytotoxic T cells in individual animals correlates with cross-protection in cattle challenged with Theileria parva. Infect Immun. 1995;63(4):1258–62.

35. Taracha ELN, Goddeeris BM, Scott JR, Morrison WI. Standardization of a technique for analysing the frequency of parasite-specific cytotoxic T lymphocyte precursors in cattle immunized with Theileria parva. Parasite Immunol. 1992 Mar;14(2):143–54.

36. Muleya W, Atuhaire DK, Mupila Z, Mbao V, Mayembe P, Kalenga S, et al. Sequence Diversity of Tp1 and Tp2 Antigens and Population Genetic Analysis of Theileria parva in Unvaccinated Cattle in Zambia’s Chongwe and Chisamba Districts. Pathogens. 2022 Jan;11(2):114.

